# Glycoengineered recombinant alpha1-antitrypsin results in comparable *in vitro* and *in vivo* activities to human plasma-derived protein

**DOI:** 10.1101/2024.03.27.587088

**Authors:** Frances Rocamora, Sanne Schoffelen, Johnny Arnsdorf, Eric A. Toth, Yunus Abdul, Thomas E. Cleveland, Sara Petersen Bjørn, Mina Ying Min Wu, Noel G. McElvaney, Bjørn Gunnar Rude Voldborg, Thomas R. Fuerst, Nathan E. Lewis

## Abstract

Alpha-1-antitrypsin (A1AT) is a multifunctional, clinically important, high value therapeutic glycoprotein that can be used for the treatment of many diseases such as alpha-1-antitrypsin deficiency, diabetes, graft-versus-host-disease, cystic fibrosis and various viral infections. Currently, the only FDA-approved treatment for A1AT disorders is intravenous augmentation therapy with human plasma-derived A1AT. In addition to its limited supply, this approach poses a risk of infection transmission, since it uses therapeutic A1AT harvested from donors. To address these issues, we sought to generate recombinant human A1AT (rhA1AT) that is chemically and biologically indistinguishable from its plasma-derived counterpart using glycoengineered Chinese Hamster Ovary (geCHO-L) cells. By deleting nine key genes that are part of the CHO glycosylation machinery and expressing the human ST6GAL1 and A1AT genes, we obtained stable, high producing geCHO-L lines that produced rhA1AT having an identical glycoprofile to plasma-derived A1AT (pdA1AT). Additionally, the rhA1AT demonstrated *in vitro* activity and *in vivo* half-life comparable to commercial pdA1AT. Thus, we anticipate that this platform will help produce human-like recombinant plasma proteins, thereby providing a more sustainable and reliable source of therapeutics that are cost-effective and better-controlled with regard to purity, clinical safety and quality.

---

Alpha-1 Antitrypsin (A1AT), the gene product derived from the human *SERPINA1* gene is a clinically important and multifunctional glycoprotein whose primary function is the inhibition of serine proteases. While its primary substrate is elastase, A1AT can neutralize many other enzymes such as proteinase-3, cathepsin G, myeloperoxidase, trypsin, and the coagulation cascade serine proteases plasmin, thrombin, urokinase, and factor Xa(de Serres & Blanco, 2014; Karatas & Bouchecareilh, 2020; Korkmaz, Horwitz, Jenne, & Gauthier, 2010). Produced primarily by liver cells, and to a lesser extent—also by macrophages, monocytes, pulmonary and intestinal cells, A1AT is one of the most abundant proteins in human plasma (1-2 g/L) and accounts for over 90% of the antiprotease activity in human serum(de Serres & Blanco, 2014; O’Brien et al., 2022). Beyond its antiprotease activity, this protein also interacts with a variety of binding partners(O’Brien et al., 2022) and exerts anti-inflammatory, immunomodulatory, tissue-protective, wound-healing effects(Lewis, 2012; O’Brien et al., 2022).

As a therapeutic protein, A1AT’s primary indication is for treating alpha-1 antitrypsin deficiency (AATD), which is one of the most common inherited disorders(de Serres & Blanco, 2014; Strnad, McElvaney, & Lomas, 2020). Patients with AATD suffer from decreased levels of functional A1AT, which predisposes them to lung and liver dysfunction and many different inflammatory conditions(de Serres & Blanco, 2014). To manage AATD, augmentation therapy can be performed—where patients are periodically infused with large amounts of A1AT to restore optimal physiological levels of the protein. Beyond AATD treatment, however, therapy using A1AT is being explored for a myriad of clinical conditions including Type I and Type II diabetes, graft-versus-host-disease, cystic fibrosis, multiple sclerosis, rheumatoid arthritis, lupus and many others(Lewis, 2012; O’Brien et al., 2022). Trials have demonstrated the positive therapeutic potential of intravenous/inhaled A1AT for respiratory viral infections, e.g., for mild to moderate COVID-19(McEvoy et al., 2021; Ritzmann et al., 2021).

Currently, commercially available A1AT that is used for clinical application is extracted from pooled plasma from human donors; thus, augmentation therapy comes with disadvantages, including its limited supply and complex purification process. Furthermore, since the protein comes from a pool of different human volunteers, there is higher batch-to-batch variation, and an added possibility of pathogen transmission from extracted plasma. Thus, many groups have sought to produce A1AT recombinantly in hosts such as yeast, bacteria, insect cells, transgenic plants and transgenic animals to increase yield and decrease cost(Karnaukhova, Ophir, & Golding, 2006); however, these systems fail to recapitulate the native human pattern of A1AT glycosylation, posing a hurdle to obtaining reliable alternative sources of a safe, affordable, and efficacious therapeutic drug. In particular, the human version of A1AT has three N-glycans that are of an A2G2S2 configuration—biantennary, afucosylated structures containing two galactose residues capped with an α-2,6-linked sialic acid(Kolarich et al., 2006; Mills et al., 2001). Recombinant forms of A1AT that lack glycosylation shows reduced stability and is rapidly cleared from the bloodstream, while the presence of non-human glycans can trigger an unwanted immune response(Karnaukhova et al., 2006; Rocamora et al., 2023). Building on prior work matching recombinant production of A1AT with human-like glycosylation(Amann et al., 2019), we have generated a cGMP-ready Chinese Hamster Ovary (CHO) cell-based platform for producing recombinant A1AT, assessed its biochemical, *in vitro*, and *in vivo* similarity to the human plasma product. In particular, our genetically-modified CHO cells yield a human-like glycosylation pattern. These glycoengineered CHO cells (geCHO-L) were subsequently made to express human A1AT with a high level of productivity and the purified recombinant product (geCHO-rhA1AT) was then compared with human plasma-derived A1AT (pdA1AT) regarding its physicochemical properties and biological activity, including *in vivo* half-life. Taken together, this study demonstrates the profound utility of the geCHO-L platform for as a reliable, more affordable system for the recombinant production of high value therapeutic proteins bearing optimal glycoprofiles.

## Glycoengineering allows for the recombinant production of human-like A1AT

Recombinantly-expressed A1AT from CHO cells results in fucosylated complex glycans, including tetra- and tri- and bi-antennary forms bearing 2,3-sialic acids(Amann et al., 2019; Lalonde et al., 2020); however, pdA1AT is almost exclusively afucosylated biantennary glycans capped with 2,6-sialic acids. To recombinantly express a fully “humanized” A1AT (rhA1AT), the producer cell line was glycoengineered to recapitulate the glycosylation profile of a native version of the protein that is derived from human plasma. To this end, GMP-ready CHO-S cells were genetically modified using CRISPR/Cas9 to induce frameshift mutations in 8 glycosytransferase genes (Mgat4a, Mgat4b, Mgat5, St3gal3, St3gal4, St3gal6, B3gnt2 and Fut8) and Sppl3—an aspartyl protease involved in cleaving various glycan-modifying enzymes(Voss et al., 2014). Mutations introduced into the abovementioned genes rendered their enzyme products nonfunctional, thus inhibiting downstream glycan branching and preventing the addition of an α-2,3-linked terminal sialic acid residues. These genetic modifications force the cell’s glycosylation machinery to generate N-glycans that are primarily afucosylated, biantennary structures with two terminal galactose residues (A2G2) (Fig. 1A)(Nagae, Yamaguchi, Taniguchi, & Kizuka, 2020; Schjoldager, Narimatsu, Joshi, & Clausen, 2020). A previous study that employed a similar multi-gene knockout approach has shown that such genetic disruptions did not negatively impact cell growth relative to wild-type CHO-S cells(Amann et al., 2019). With the exogenous expression of a human β-galactoside α-2,6-sialyltransferase 1 (hST6GAL1), the CHO-S cells can cap this A2G2 structure with an α2,6-linked sialic acid, thereby producing the dominant N-glycoform that is found in human A1AT (A2G2S2)(Clerc et al., 2016; Kolarich et al., 2006; Mills et al., 2001). After FACS-sorting and clonal expansion of 9xKO cells that also stably express hST6GAL1 we obtained what are here referred to as geCHO-L cells. These were then transfected with an expression plasmid containing the coding sequence for human A1AT (Fig. 1A), and we identified clones that stably produce up to 1.6 g/L of pure recombinant human A1AT (geCHO-rhA1AT). The geCHO-rhA1AT was then purified from the supernatant of a high producing clone and subjected to chemical and biological characterization in comparison with commercially available pdA1AT. Using size exclusion chromatography coupled to multiple angle light scattering (SEC-MALS) and dynamic light scattering (DLS), the size and composition of both A1AT species were investigated. Comparative analysis showed that commercially available pdA1AT (purchased from Athens Research & Technology), and purified geCHO-rhA1AT have a comparable molecular weight (*p = 0*.*43*) and hydrodynamic radius (*p = 0*.*46*) (Fig. 1B, Table 2).

**Figure 1.**
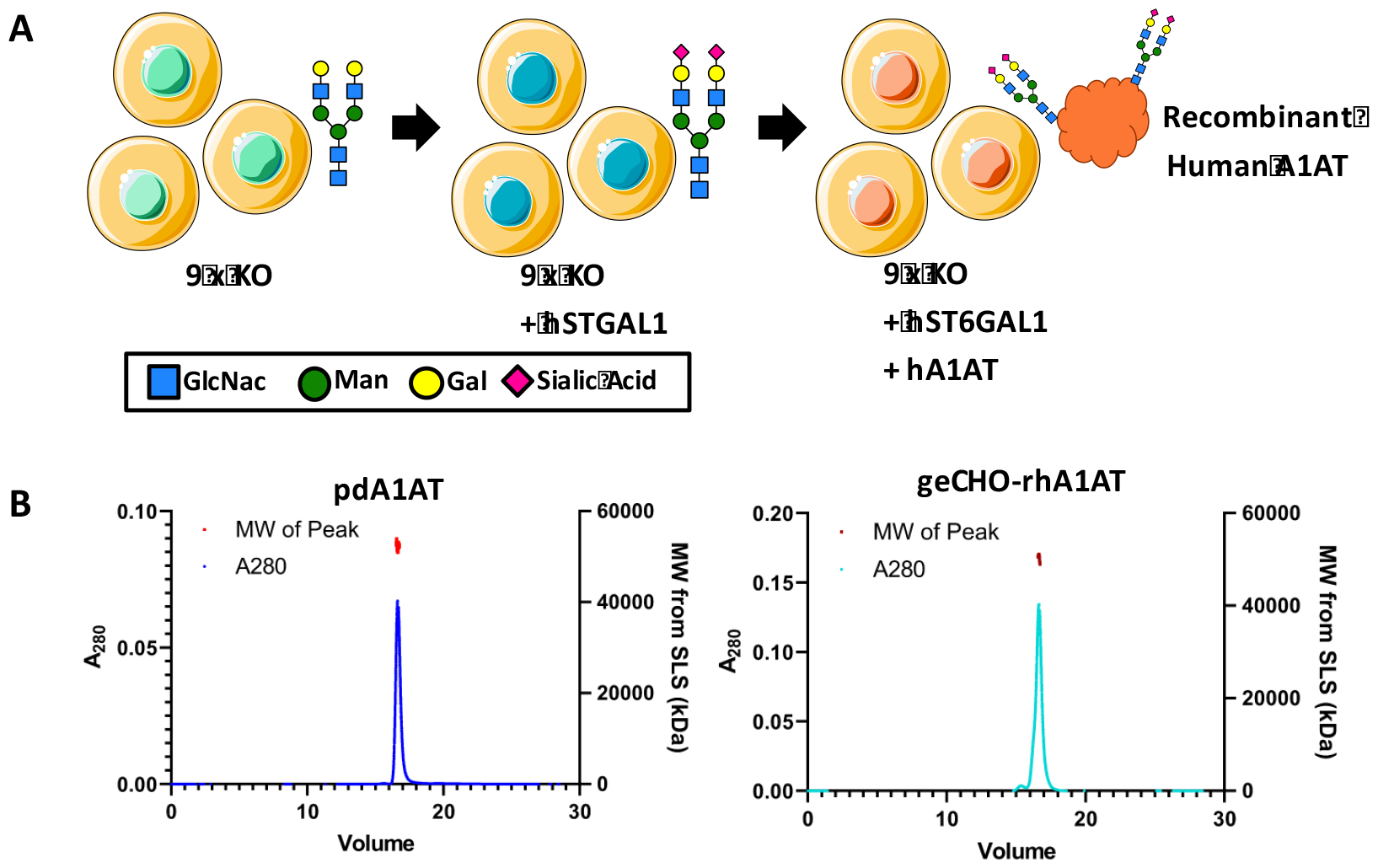
Production of rhA1AT Using Glycoengineering. (A) To produce A1AT with a human-like glycan profile, a CHO parent clone whereby 9 genes involved in protein glycosylation (Mgat4a, Mgat4b, Mgat5, St3gal3, St3gal4, St3gal6, B3gnt2, Fut8 and Sppl3) have been knocked out were made to express human ST6GAL1 and human A1AT via random integration. (B) The molecular size of plasma-derived and recombinant A1AT was also analyzed using SEC-MALS and DLS. The chromatographs of each protein are shown as blue (pdA1AT) or cyan (geCHO-rhA1AT) lines. The MALS scattering sizes between the peak half-maxima are shown as red (pdA1AT) or maroon (geCHO-rhA1AT) points.

## geCHO-rhA1AT cells have similar N-glycan profile as pdA1AT

Using LC-MS, we evaluated the glycosylation of the rhA1AT and pdA1AT. N-glycan analysis revealed that the purified geCHO-rhA1AT and pdA1AT also bear nearly identical glycan profiles (Fig. 2). While changes in the glycan profile of A1AT has been observed among human subjects afflicted pulmonary disease(Liang et al., 2015; Rho, Roehrl, & Wang, 2009; Wen et al., 2012), the most abundant isoform of A1AT identified in healthy patients is a biantennary, disialylated (with an α-2,6-linked sialic acid), afucosylated glycan structure(Clerc et al., 2016; Kolarich et al., 2006; Mills et al., 2001). As expected, this N-glycan represents the largest peak (∼85%) detected through LC-MS in the commercially available pdA1AT (Fig. 2A, Table 1). Our geCHO-L cells produced this exact structure in geCHO-rhA1AT (Fig. 2B). An additional benefit of this cell line is that it not only produces the desired glycoform of the recombinant protein of interest, but that by using a clonal, glycoengineered parent for the production of A1AT, the proteins produced also show a considerably higher level of glycoprofile homogeneity in the purified product relative to recombinant A1AT expressed in a wild-type CHO-S parent(Amann et al., 2019; Lee et al., 2013) (Fig. 2C). N-glycan profiling shows that rhA1AT bears the A2G2S2 form almost exclusively (∼72%) with the largest peak representing the mature form of this glycan species, while the smaller peaks represent either precursors or subspecies of A2G2S2 (Fig. 2B, Table 1); meanwhile the desired glycoform is undetected when A1AT is produced in the wild-type CHO-S cells (Fig. 2C, Table 1).

**Table 1.** Glycan Structures Identified using LC-MS.

| **Protein** | **Glycan** | **% Peak Area** |
| --- | --- | --- |
| **pdA1AT** | A2G2 | 0.2 |
|  | A2G2S2* | 4.9 |
|  | A2G2S2 | 84.1 |
|  | A2G2S2-F | 4.7 |
|  | A3G3S3 | 3.2 |
|  | A3G3S3-F | 2.8 |
| **geCHO-rhA1AT** | M5 | 1.5 |
|  | A1 | 2.3 |
|  | A2 | 7 |
|  | A1G1S1 | 1.7 |
|  | A2G2 | 0.3 |
|  | A2G1S1 | 14.1 |
|  | A2G2S1 | 0.7 |
|  | A2G2S2 | 72.3 |
| **rhA1AT from**<br>**WT CHO-S** | A1F | 1.9 |
|  | A2F | 3.8 |
|  | A2FG2 | 10.3 |
|  | A2FG2S1 | 13.8 |
|  | A2FG2S2 | 35.5 |
|  | A2G2S1 | 2.5 |
|  | A3FG3S2 | 1.6 |
|  | A3G3S3 | 6.9 |
|  | A4FG4S3 | 1.1 |
|  | A4FG4S4 | 1.9 |
|  | M3 | 0.5 |
|  | M5 | 4.4 |
|  | M6 | 4.8 |
|  | M7 | 3.8 |
|  | M8 | 3.9 |
|  | M9 | 3.1 |

**Figure 2.**
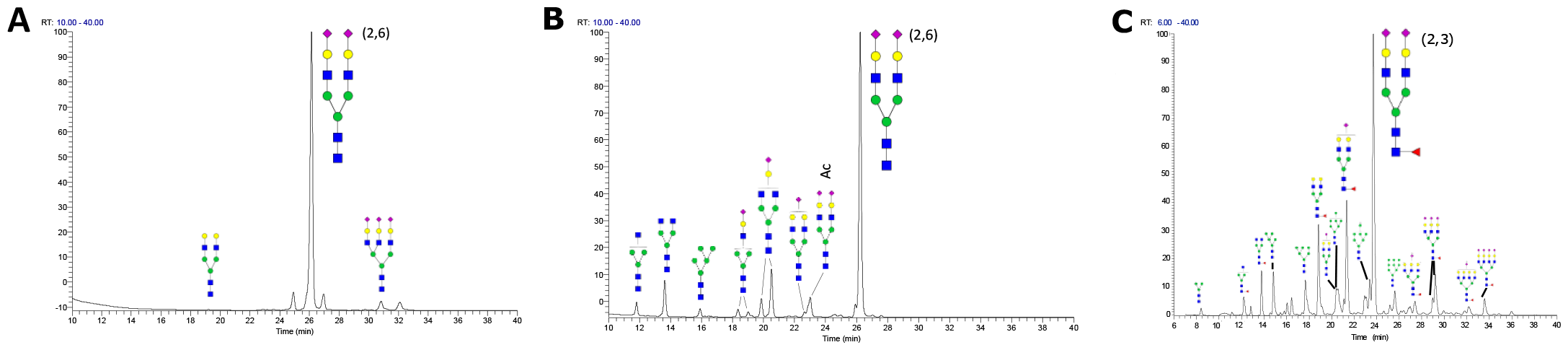
Glycan Analysis of Plasma-derived and recombinant A1AT. N-glycan analysis of (A)pdA1AT, (B)geCHO-rhA1AT, and (C) wild-type CHO-derived recombinant A1AT was performed using LC-MS.

## geCHO-rhA1AT exhibits both activity and half-life that are indistinguishable from pdA1AT in terms of activity and half-life

Finally, we compared the biological activity, and the pharmacokinetic profile of geCHO-rhA1AT and pdA1AT. To test the ability of both proteins to inhibit serine proteases, an *in vitro* assay that quantifies elastase activity was carried out. Indeed, we found comparable elastase inhibition for recombinant (IC_50_ = 69.1 nM) and plasma-derived (IC_50_ = 67.8 nM) proteins (Fig. 3A, Table 2). This is unsurprising given the physicochemical similarity between pdA1AT and geCHO-rhA1AT. Although it has been observed that losses of N-glycosylation of A1AT impacts protein stability and enzymatic activity, smaller changes in glycan structure are more likely to impact blood half-life and clearance(Karnaukhova et al., 2006).

**Table 2.** Biochemical characteristics of pdA1AT and geCHO-rhA1AT.

| Feature | pdA1AT | geCHO-rhA1AT |
| --- | --- | --- |
| Molecular weight (kDa) | 51.7 ± 1.5 | 50.0 ± 1.2 |
| Hydrodynamic radius (nm) | 3.5 ± 0.1 | 3.4 ± 0.2 |
| Elastase IC<sub>50</sub> (nM) | 67.8 ± 4.8 | 69.1 ± 0.9 |
| t<sub>1/2</sub> (hrs) | 5.0 ± 0.6 | 6.0 ± 0.4 |

**Figure 3.**
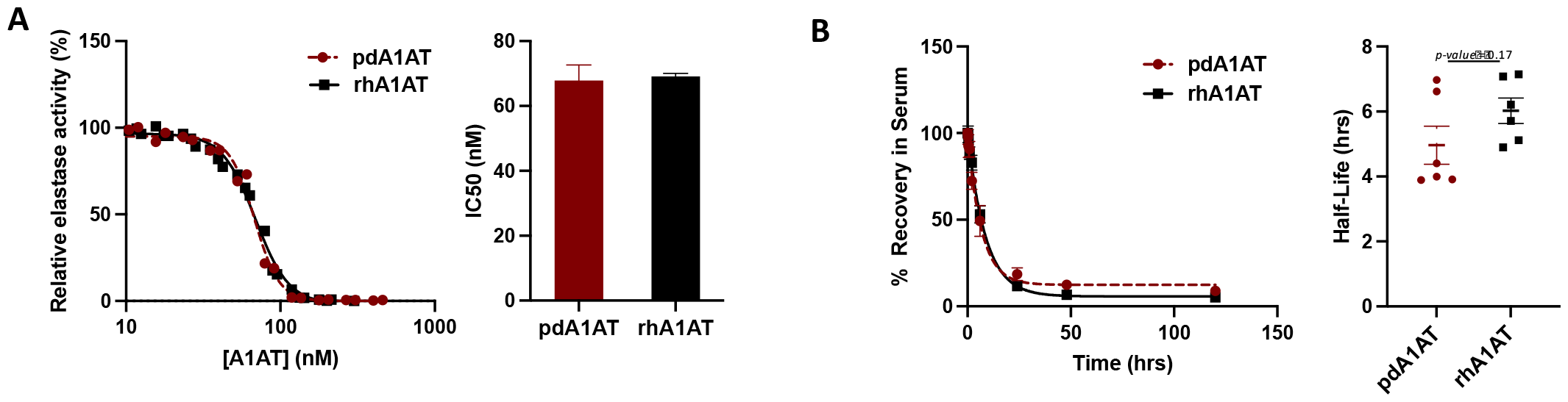
Comparing *in vitro* activity and *in vivo* half-life of rhA1AT and pdA1AT. (A) *In vitro* inhibitory activity of purified rhA1AT against elastase was evaluated and compared with that of commercial *pd*A1AT. Data shown represents at least two experimental replicates (mean ± SEM). (B) Half-life of pdA1AT and rhA1AT was measured in SD rats after a single-dose, intravenous injection at a concentration of 150 µg/kg. Blood was collected at 5mins, 15 mins, 30 mins, 1 hour, 2 hours, 6 hours, 24 hours, 48 hours, and 120 hours post-injection, and levels of either protein remaining in the serum was measured using sandwich ELISA. Figure represents data from 6 animals per group, with each group receiving an injection of either pdA1AT or geCHO-rhA1AT (mean ± SEM).

Next, we evaluated the *in vivo* clearance rate of each version of A1AT in male and female Sprague-Dawley rats by measuring the relative abundance of either pdA1AT or geCHO-rhA1AT remaining in plasma over time from a single tail vein injection of 150 µg/kg. Our results demonstrate that there is no statistically significant difference between the circulatory half-life of pdA1AT (t_1/2_ = 5 hrs) and geCHO-rhA1AT (t_1/2_ = 6 hrs) (Fig. 3B, Table 2) (*p=0*.*17*). This finding is especially important because of the profound impact that N-glycosylation has on the stability and serum longevity of many therapeutic proteins, including A1AT(Rocamora et al., 2023). Absent or aberrant glycosylation previously stymied past attempts to recombinantly express human A1AT in non-human host systems, wherein the glycoforms obtained had inferior stability and clearance profiles compared to the native pdA1AT(Karnaukhova et al., 2006; Rocamora et al., 2023). Furthermore, it has been shown that proper glycosylation is essential to the capacity of A1AT to bind IL-8 and regulate neutrophil chemotaxis(Bergin et al., 2010) and changes in the level of sialylation can significantly impact the anti-inflammatory activity of the protein(McCarthy et al., 2018). While a previous study(Amann et al., 2019) also demonstrated that recombinant A1AT with a human-like glycoprofile could be produced by manipulating the glycosylation machinery of CHO cells, this is the first report corroborating a near-identical pharmacokinetic profile between the plasma-derived protein and the recombinantly expressed, glycoengineered product.

By recombinantly expressing the human coding sequence of A1AT in geCHO cells that have been modified to mimic the same glycoprofile as that of pdA1AT, we generated a GMP-ready clone that secretes >1g/L of recombinant protein. This level of productivity is similar to what has been achieved from previous attempts to produce A1AT in CHO cells(Amann et al., 2019; Lalonde et al., 2020; Lee et al., 2013), but this time with the added advantage of generating a highly homogenous batch of rhA1AT that contains identical N-glycans as its human counterpart, pdA1AT. Importantly, we show that the rhA1AT obtained from geCHO-L is indistinguishable from pdA1AT in terms of elastase inhibitory activity and elimination half-life, making it a highly promising alternative source of therapeutic A1AT for clinical use in the future. Given the comprehensive and long-standing application of CHO-based expression systems in the pharmaceutical industry, the platform described here offers many great advantages for producing therapeutic A1AT regarding safety, efficiency and cost relative to the current model of harvesting the protein from human plasma.

Beyond the ability to perfectly capture human-like glycan structures, glycoengineering CHO cells can also be utilized to produce therapeutic proteins having optimized glycoprofiles that impart improved binding, heightened specificity or better pharmacokinetic profiles. Modifying the native glycosylation profiles of many biologics encompassing monoclonal antibodies, protein vaccines and hormones have resulted in better therapeutic outcomes(Chen et al., 2023; Lusvarghi et al., 2023; Newby, Allen, & Crispin, 2023; Rocamora et al., 2023), and tools are emerging to enable more predictive glycoengineering of therapeutics.(Spahn, Hansen, Kol, Voldborg, & Lewis, 2017; Spahn & Lewis, 2014; Yang et al., 2015) In the case of A1AT, different diseases states of inflammation and malignancy have been shown to be associated with alternative glycoforms of the protein suggesting that variations in glycosylation could play a role in tuning the protein’s immunomodulatory properties(McCarthy et al., 2014). In the future, we therefore hope to expand this study by utilizing the geCHO platform to produce and investigate new, pharmacologically superior glycovariants of A1AT.

## Materials and Methods

### Generation of the CHO-S clone stably expressing rhA1AT with a glycoprofile similar to the human plasma derived A1AT

CHO-S cells were cultured and handled according to the Gibco guidelines. First, a multiplex 9X knockout clone, with frame-shift mutations in eight glycosyltransferase genes (Mgat4a, Mgat4b, Mgat5, St3gal3, St3gal4, St3gal6, B3gnt2 and Fut8) and the Sppl3 gene, was generated from the CHO-S parental cell line using the CRISPR/Cas9 gene editing technology described in detail previously by Amann et al(Amann et al., 2019). Secondly, this clone was used to generate a stable pool harboring the human glycosyltransferase gene ST6GAL1 via transfection with a plasmid containing the coding sequence of human beta-galactoside alpha-2,6-sialyltransferase 1 isoform a (hSTGAL1) (GenBank NP_001340845.1) and expressed from a composite promoter mCMV-hEF1-HTLV (InvivoGen, San Diego, CA) in a derivative of pcDNA3.1(+) (Life Technologies, Carlsbad, CA).

Cells from this pool were FACS single cell sorted and lectin stained according to the procedure described previously(Amann et al., 2019), and a clone verified to have a high and stable expression of human ST6GAL1 was chosen as a parental cell line for the next and final round of transfection. By random integration, a clonal 9xKO-hSTGAL1-expressing CHO parent was made to express human A1AT via transfection with a plasmid containing the coding sequence of human alpha-1-antitrypsin (hA1AT) (GenBank NP_000286.3) under a composite promoter mCMV-hEF1-HTLV (InvivoGen, San Diego, CA) in a pcDNA3.1/Zeo(+) (Life Technologies, Carlsbad, CA) backbone. To confirm production of human A1AT, the transfected population were stained with FITC conjugated anti-A1AT antibodies (abcam ab19170) and cells with the strongest signal were sorted using FACS. Titer and confluence data were obtained from 1920 clones (5 x MD384 plates) by Octet (Satorius) and Celigo (Nexcelom) analyses. Clone 1-C2 was chosen for subsequent batch cultivation and rhA1AT protein purification.

Coding sequences for hSTGAL1 and hA1AT were both codon-optimized for expression in Chinese Hamster (*Cricetulus griseus*) cells and synthesized by Geneart, Regensburg, Germany.

### Expression of rhA1AT for PK studies

Cryopreserved clone 1-C2 cells were thawed and cultured for 3 passages before inoculation of a 2L Corning shake flask, containing 500 mL complete CD-CHO medium, with 1x10E6 viable cells/mL. Cells were cultured for 7 days with shaking at 120 rpm in a humidified CO2-incubator supplied with 5% CO2. Supernatant was harvested by first centrifugation at 300g for 5 minutes and secondly cleared by another centrifugation at 1000g for 10 minutes before protein purification was initiated.

### Protein purification

Cell culture material containing rhA1AT was centrifuged at 13,000g for 10 minutes at 4°C. Supernatant (460 mL) was collected and desalted by tangential flow filtration (TFF) using a Centramate T-Series Cassette (Pall, DC010T02) mounted in a Centramate LV holder (Pall). First, the sample was up-concentrated to a final volume of 60 mL. Next, one diafiltration step was performed using 250 mL of 25 mM Bis-Tris buffer, pH 5.92. Following the diafiltration step, the product was collected and the TFF membrane was rinsed twice with 50 and 25 mL buffer, respectively, in order to recover as much product as possible.

*rh*A1AT was purified on an ÄKTA Pure protein purification system following a two-step anion exchange chromatography procedure. Both steps were performed using Q-Sepharose columns (Cytiva), the first step with 25 mM Bis-Tris buffer (pH 5.92) and the second step with 10 mM Na_2_HPO_4_, 2 mM KH_2_PO_4_ (pH 8.0). In the first step, the wash step contained 50 mM NaCl, while the protein was eluted with 150 mM NaCl. In the second step, the column was washed with 70 mM NaCl and the protein was eluted over a linear gradient from 70 to 200 mM NaCl. Fractions containing the protein of interest were concentrated on an Amicon® centrifugal filter device (10 MWCO) and temporarily stored at -80 °C. As a final polishing step, the protein was thawed and loaded onto a Superdex200 increase 10/300 column (Cytiva) using dPBS as eluent. Fractions of interest were pooled, concentrated to <4 mL and loaded onto a 0.5 mL Pierce high-capacity endotoxin removal spin column. Next, the concentration was adjusted to 1.15 mg/mL by dilution in dPBS. The solution was sterile-filtered, and stored at -80 °C.

### Size exclusion chromatography coupled to multiple angle light scattering (SEC-MALS) and dynamic light scattering (DLS)

For SEC-MALS, a Vanquish Flex UHPLC system (Thermo Fisher Scientific^*^) was coupled to Dawn HELEOS-II MALS (Wyatt) and Optilab T-rEX refractive index (Wyatt) detectors. One detector position from the MALS instrument was replaced with an optical fiber coupled to a DynaPro NanoStar DLS instrument (Wyatt), allowing for simultaneous UV, MALS, DLS and RI measurements. Separations were performed using either an SRT SEC-300 7.8x50mm column (Sepax) or a Superose6 Increase 10/300 GL column (Cytiva) equilibrated in PBS with a flow rate of 0.3 mL/min and sample injection volumes of 25 μL. Molar mass (z-weighted) was determined using the standard workflow within the software ASTRA 7.3.2.17 (Wyatt) using refractive index as a concentration source. Hydrodynamic radius from DLS (z-weighted) was also determined using ASTRA. Molar mass and hydronamic radius measurements were performed 4 times for each of the two proteins using the same instrument over a 6-month period, and are reported as mean ± s.e.m. Values for the two proteins were compared using an independent-samples t-test (two tailed).

### Glycan analysis

N-glycans were derivatized with GlycoWorks RapiFluor-MS N-Glycan Kit (Waters, Milford, MA) according to the manufacturer’s instruction. Briefly, 12 μg purified protein was used for each sample. Labeled N-Glycans were analyzed by LC-MS as described previously(Grav et al., 2015). Separation gradient from 30% to 43% 50 mM ammonium formate buffer and MS were run in positive mode. Amount of N-Glycan was measured by integrating the peaks with Thermo Xcalibur software (Thermo Fisher Scientific, Waltham, MA) giving the normalized, relative amount of the glycans.

### Enzyme Activity Assay

Anti-elastase activity was determined using the EnzChek™ Elastase Assay kit (Invitrogen™). A 1.5-fold dilution series of A1AT was prepared, with 460 nM of A1AT as the highest inhibitor concentration and with 10 concentrations in total (minimum of 120 µL per dilution). In a black, clear-bottom 96-well plate, first 100 µL of elastase working solution (0.5 U/mL) was added to the relevant number of wells (10 + 1 wells per A1AT variant, in duplicate). Next, 50 µL of each A1AT dilution was added to 10 consecutive wells containing the elastase working solution. As reference, 50 µL reaction buffer instead of A1AT solution was added to the 11th well. Using a multichannel pipette, 50 µL of substrate solution (0.1 mg/mL) was added to all 11 wells at the same time. Solutions were mixed by pipetting up and down. For background correction, the 12th well in a row was filled with 150 µL assay buffer and 50 µL substrate solution. The plate was incubated at room temperature protected from light for 50 min. Fluorescence intensity was measured in a fluorescence microplate reader (BioTek Synergy Mx), λ_ex_ set at 485/9.0 nm and λ_em_ at 530/13.5 nm, 10 reads per well, gain at 80.

Data was processed by first subtracting the averaged background fluorescence from averaged intensities measured for each A1AT concentration and the reference. Next, the relative elastase activity detected in wells containing A1AT was normalized to the activity detected in the reference well. Relative elastase activity values were plotted against A1AT concentration in GraphPad Prism. IC_50_-values were derived by non-linear regression using the “[Inhibitor] vs normalized response – variable slope” as model.

### Ethics Statement

This IACUC-approved study (Approval Number NLS-650) was conducted in compliance with the following: Animal Welfare Act Regulations (Title 9 of the Code of Federal Regulations); US Public Health Service Office of Laboratory Animal Welfare (OLAW) Policy on Humane Care and Use of Laboratory Animals; the Guide for the Care and Use of Laboratory Animals (Institute of Laboratory Animal Resources, Commission on Life Sciences, National Research Council, 1996)(1996); and AAALAC accreditation.

### Animal Study Design

Male and female Sprague Dawley rats (8 weeks old) weighing 226 ± 23 g were used for this study. Two groups each consisting of three males and three females were injected intravenously via tail vein with 1.5 mL of either rhA1AT or pdA1AT at a dose of 150 µg/kg. Blood was collected from each animal into heparin tubes at ten time points: right before injection, and at 5minutes, 15 minutes, 30 minutes, 1 hour, 2 hours, 6 hours, 24 hours, 48 hours, and 120 hours post-injection for evaluation of A1AT abundance.

### Detection of A1AT levels in plasma

Collected blood extracted from each animal was processed into plasma and levels of rhA1AT or pdA1AT were measured via sandwich ELISA using a matched antibody pair specific for human A1AT (Novus Biologicals NBP2-79565) and following the manufacturer’s instructions. The detection antibody is conjugated to horseradish peroxidase, and the chromogenic substrate used for detection is TMB (Sigma). Before quantification, plasma samples were diluted in 1:180 in a sample buffer made of 0.1% BSA in 0.05% TBST. For signal detection, absorbance at 450 nM was measured using a microplate reader (BioTek Synergy Mx). The relative abundance of either pdA1AT or rhA1AT was normalized to the signal obtained from the first timepoint (5 mins post-injection) which consistently showed the highest level of protein recovered. Half-lives from each animal was calculated by fitting the data to a one-phase exponential decay curve using GraphPad Prism.

## Acknowledgements

This work was supported in part by generous funding from the Novo Nordisk Foundation (NNF20SA0066621) and NHLBI (R41 HL164260)

## Footnotes

* Certain commercial products or company names are identified here to describe our study adequately. Such identification is not intended to imply recommendation or endorsement by the National Institute of Standards and Technology, nor is it intended to imply that the products or names identified are necessarily the best available for the purpose.

